# Fixing Hybrid Rice: >99% Efficient Apomixis with Near-Normal Seed Set

**DOI:** 10.1101/2025.10.16.682968

**Authors:** Wen-Qiang Chen, Chaolei Liu, Hongwei Lu, Shenlin Yang, Jinxin Ma, Yong Huang, Jie Xiong, Derun Huang, Zhihuan Tao, Jianrong Lin, Kejian Wang

## Abstract

Apomixis, a form of clonal seed reproduction, offers a transformative approach to agriculture by enabling the stable fixation of hybrid vigor and elite heterozygosity across generations. However, the practical implementation of synthetic apomixis has been hindered by highly variable clonal efficiency or significant yield penalties. In this study, through integrated transcriptomic analyses, we identified a sperm-specific transcription factor in rice that likely functions as a key initiator of embryogenesis. Ectopic expression of this factor in egg cells effectively induces parthenogenesis and produces haploid progeny. When combined with clonal gametogenesis, this system achieved nearly complete synthetic apomixis, with clonal seed production rates exceeding 99% across all derived hybrid rice lines. Moreover, we generated apomictic hybrid lines that not only consistently produced over 99% clonal seeds but also exhibited seed yields comparable to conventional F_1_ hybrids. These findings establish a scalable and agriculturally viable platform for synthetic apomixis, achieving stable fixation of heterosis through clonal seeds and paving the way for the commercial deployment of self-perpetuating hybrid crops.

## INTRODUCTION

Heterosis, or hybrid vigor, refers to the superior performance of hybrid offspring relative to their inbred parents, and has been extensively exploited in major crops such as rice and maize, resulting in substantial improvements in global agricultural productivity.^1^ For instance, hybrid rice varieties typically outperform their inbred counterparts by 20–30% in yield.^2,3^ However, due to sexual reproduction, hybrids undergo genetic segregation in subsequent generations, leading to variable phenotypes and a loss of heterotic traits. As a result, hybrid seeds must be regenerated each season via technically demanding and costly crossing procedures.^4,5^ Engineering seed-based clonal reproduction—known as synthetic apomixis—offers a promising strategy to permanently fix heterosis and preserve elite heterozygous genotypes across generations.^6-10^ To realize this goal at scale, it is essential to achieve clonal seed production with complete or near-complete penetrance, while maintaining normal seed set and yield potential.^11-14^

Our previous studies have developed genome-editing strategies that combine clonal gametogenesis (*Mitosis instead of Meiosis, MiMe*)^15,16^ with mutations in haploid induction genes specifically expressed in sperm or pollen, such as *OsMTL* and *OsPLA*α2 in rice, and *AtDMP8/9* in *Arabidopsis*.^17-19^ Although this genome-editing-only approach is independent of T-DNA insertion, its practical utility is limited by low clonal efficiency. A more effective strategy is to integrate *MiMe* with the induction of parthenogenesis—the autonomous development of egg cells into embryos without fertilization.^12-14^ Ectopic expression of transcription regulators in egg cells, including endogenous transcription factors such as *BABY BOOM 1* (*OsBBM1*), *OsBBM4*, and *WUSCHEL* (*OsWUS*),^20-22^ as well as heterologous regulators *PARTHENOGENESIS* (*PpPAR* and *ToPAR*),^23-26^ has been shown to trigger parthenogenesis in rice. While *OsBBM1*-based systems can achieve clonal seed production rates of 11%–29%,^20^ optimized versions have reported efficiencies exceeding 95%.^27-29^ However, clonal efficiencies still vary widely among lines, and most lines exhibit 0%–90% clonal efficiency and are frequently accompanied by severely reduced seed set.^20,27-29^ Similar challenges persist with *OsBBM4, OsWUS, PpPAR*, and *ToPAR*, where fertility may be improved but clonal efficiency remains suboptimal.^21-24^

A long-standing hypothesis posits that egg cells remain mitotically quiescent and require sperm-derived factors to initiate cell division and embryogenesis.^30-34^ Indeed, the egg cell is transcriptionally poised and rapidly activates transcriptional programs upon fertilization.^35-39^ Interestingly, all endogenous rice genes known to induce parthenogenesis—including *OsBBM1, OsBBM4*, and *OsWUS*—encode broadly expressed transcription factors.^20-22^ These genes are broadly active in multiple tissues—including roots, shoots, panicles, endosperm, and embryos—and loss-of-function mutations in these genes impairs embryogenesis, endosperm development, seed germination, tiller formation, and overall plant growth.^20,40-42^ These observations raise a compelling question: do specialized transcription factors exist that are specifically expressed in sperm cells and can trigger the transition of the egg cell into an embryo? Such factors might more precisely recapitulate the natural mechanisms of embryogenesis, providing a refined and biologically aligned approach to parthenogenesis induction. Their identification and application could substantially advance the establishment of robust synthetic apomixis systems for crop improvement in agricultural practice.

Here, we identified a sperm cell–specific transcription factor in rice through integrative analysis of tissue-specific transcriptomics, developmental time-series data, and gene co-expression networks. Ectopic expression of this gene in egg cells efficiently induced parthenogenesis and haploid formation. When combined with clonal gametogenesis, this strategy achieved near-complete clonal seed production across all tested hybrid lines. Remarkably, we also obtained hybrid rice lines that exhibited stable inheritance, normal or near-normal seed set, and >99% clonal reproduction. These findings highlight the substantial potential of this sperm cell-specific embryogenesis trigger and the established synthetic apomixis system as an agronomically viable strategy to stably fix heterosis through seeds.

## RESULTS

### Identification of sperm-specific and delivered transcription factors

To identify potential sperm-specific transcription factors that could trigger embryogenesis, we performed a systematic transcriptome-wide analysis employing 8,796 publicly available rice RNA-seq datasets.^42^ To further eliminate the batch effect in large-scale RNA-seq data, we transformed the FPKM matrix in the database into a TPM expression matrix (Table S1). From this matrix, we initially identified 729 sperm-specific genes through a tissue-specific gene enrichment analysis (Figure S1). Sperm-delivered factors are expected to be highly expressed in sperm cells, detectable in zygotes, and absent or minimally expressed in egg cells, as their presence in zygotes originates from sperm-derived transcripts. To pinpoint genes with expression patterns indicative of sperm delivery, we further performed time-series analysis across sperm cells, egg cells, zygotes, and embryos (Figure S2). Among ten identified clusters, two (Cluster 1 and Cluster 2) aligned with the expected profile, making these genes as strong candidates for triggering embryogenesis, and were selected for further analysis (Figures 1A and S2). Among 170 genes in these two clusters, five were predicted to encode transcription factors. To further elucidate the functional roles of these candidate transcription factor genes, we constructed a co-expression network using the normalized expression matrix (Table S1). In this network, genes exhibiting similar expression patterns are grouped together, reflecting their function in tightly linked biological processes. The result showed that four out of the five transcription factor genes co-localized within a single regulatory module (Figure 1B). Notably, one gene displayed the highest connectivity and average co-expression weight within this module, suggesting it functions as a major regulatory hub (Figure 1B). We named it *HUAXU* (*华胥*)—remarkable for its ability to induce parthenogenesis (see below)—after a mythological female figure who conceived Fuxi (伏羲) and Nvwa (女娲) upon stepping into a divine footprint according to ancient Chinese texts. This new nomenclature not only facilitates reference but symbolically captures the gene’s function in inducing embryogenesis without paternal DNA contribution.

**Figure 1.**
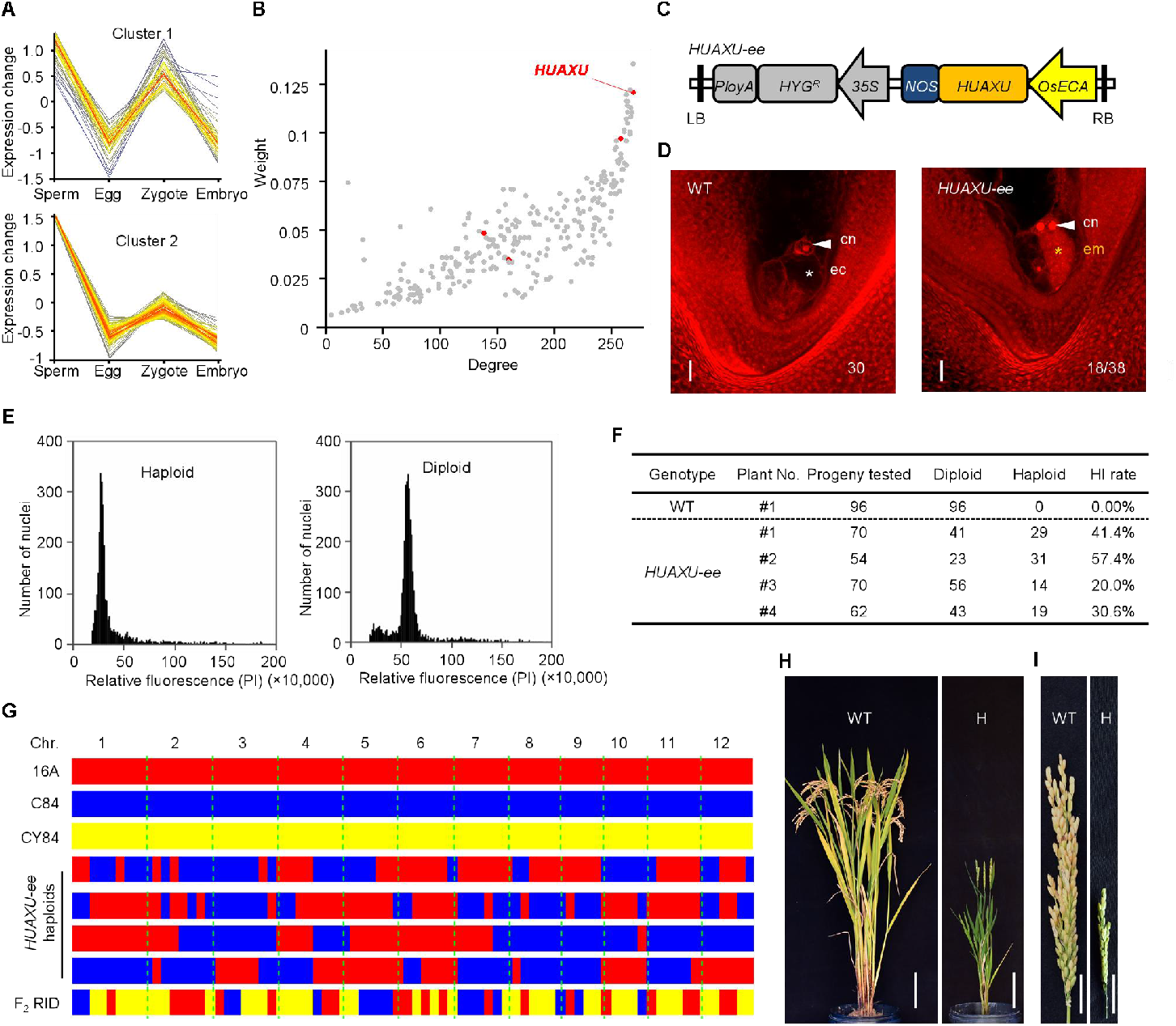
Ectopic expression of *HUAXU* in egg cells efficiently induces parthenogenesis and haploid progeny formation. (A) Cluster analysis showing expression patterns characteristic of sperm-delivered zygotic factors, based on time-series data from sperm cells, egg cells, zygotes, and embryos. (B) Gene co-expression weights within the cluster containing four sperm-specific candidate transcription factors. These four genes are highlighted in red. Among them, *HUAXU* exhibits the highest weight, indicating its role as a major hub gene. (C) Schematic of the binary vector construct (*HUAXU-ee*) designed for egg cell-specific ectopic expression of *HUAXU. HYG*^*R*^, hygromycin resistance gene. (D) Parthenogenetic embryos (orange asterisks) induced by egg cell-specific *HUAXU* expression in ovaries of an emasculated *HUAXU-ee* plant at 3 days after emasculation (DAE; n = 18/38). In wild-type (WT) ovaries, no multicellular embryo structures are observed, with only egg cells present (n = 30). In the absence of fertilization, no endosperm development is observed, and only central cell polar nuclei (white arrows) are detected in both WT and *HUAXU-ee* ovaries. Embryos (em; orange asterisks), central nuclei (cn; white arrows), and egg cells (ec; white asterisks) are indicated. Scale bars, 20 μm. (E) Flow cytometry analysis of *HUAXU-ee* progeny showing both haploid (left) and diploid (right) individuals. PI, propidium iodide. (F) Ploidy analysis of progeny from WT and *HUAXU-ee* T_0_ lines. Ploidy was assessed by flow cytometry. HI rate represents haploid induction rate. (G) Whole-genome sequencing of haploid progeny derived from *HUAXU-ee* T_0_ lines. SNPs specific to the 16A allele are shown in red, C84 in blue, and mixed alleles in yellow. Haploid progeny display genome-wide homozygosity from either 16A or C84, with no heterozygous regions. By contrast, an F_2_ recombined inbred diploid (RID) of CY84 shows recombination and partial heterozygosity. (H-I) Morphology of plants (H) and panicles (I) of WT (left) and haploids derived from *HUAXU-ee* (right). Scale bars, 20 cm (H) and 2 cm (I).

### Egg cell-specific expression of *HUAXU* triggers parthenogenetic embryogenesis in hybrid rice

The sperm-delivered transcription factors have been proposed to act as potential and critical initiators of embryogenesis following sperm-egg fusion.^34-37^ To evaluate whether *HUAXU* could be engineered to induce parthenogenesis, we constructed a binary vector, *HUAXU-ee*, designed to drive ectopic expression of *HUAXU* specifically in egg cells under the control of the rice *OsECA* promoter (Figure 1C), which has been previously validated for egg cell-specific activity.^44^ We then introduced the *HUAXU-ee* construct into the hybrid rice cultivar Chunyou 84 (CY84, wild type, WT) via *Agrobacterium*-mediated transformation of callus tissues, generating four independent lines harboring the *HUAXU-ee* transgene expression cassette (Figure S3). To assess whether *HUAXU* expression in egg cells could initiate autonomous embryogenesis without fertilization, we cultivated *HUAXU-ee* T_0_ plants under field conditions, collected pistils from *HUAXU-ee* at 3 days after emasculation (3 DAE), and detected them via Eosin B staining.^45^ The results revealed that pistils from WT plants consistently contained a single egg cell and two undivided central nuclei, with no evidence of embryogenesis or fertilization (n = 30; Figure 1D left). In contrast, multicellular embryo-like structures were observed exclusively in 18 out of 38 *HUAXU-ee* pistils, accompanied by unfertilized central cells—hallmarks of autonomous embryo development (Figure 1D right). These results demonstrate that egg cell-specific expression of *HUAXU* can efficiently induce parthenogenesis and initiate the embryogenesis of egg cells in the absence of fertilization.

### Egg cell-specific expression of *HUAXU* efficiently generates haploid progeny

Engineered parthenogenesis in sexual plants theoretically enables the production of haploid embryos and progeny, representing a key strategy for haploid induction in plant breeding.^46-47^ Before assessing whether egg cell-specific expression of *HUAXU* can trigger haploid formation, we first investigated the field performance of *HUAXU-ee* T_0_ plants. We observed that the *HUAXU-e*e T_0_ plants displayed similar vegetative and reproductive growth to WT controls (Figures S4A and S4B). However, seed set varied among the four lines, ranging from 50.3 ± 4.2% to 74.6 ± 4.3%, compared to 74.3 ± 2.5% in WT plants (Figures S4B and S4C). We then germinated seeds from the four self-pollinated *HUAXU-ee* T_0_ lines and cultured the resulting T_1_ plants. We determined the nuclear ploidy of the T_1_ progeny generated from *HUAXU-ee* lines via flow cytometry analysis using leaf tissue. The results revealed that among 54 to 70 individual plants analyzed per line, 20.0% to 57.4% of the progeny generated from *HUAXU-ee* lines were haploids. In contrast, all offspring from WT plants were determined as diploids (n = 96; Figures 1E, 1F, and S5). Another characteristic of haploids is their completely homozygous genotypes across the entire genome.^48^ To confirm the haploid identity of *HUAXU-ee* progeny, we used 12 insertion-deletion (InDel) markers, polymorphic between the parental lines of CY84 F_1_ hybrids and spanning all 12 chromosomes, to determine their genotypes (Table S2). The results revealed that a recombined inbred diploid (RID) of the hybrid CY84 showed partial heterozygosity, whereas all identified haploids exhibited homozygous genotypes identical to either the maternal or paternal genomes of hybrid CY84 at all 12 loci (Figure S6). To further validate the homozygosity across the entire genome, we resequenced the haploids, achieving an average coverage of 20×. The results demonstrated that, in contrast to the RID control plants, which exhibited both homozygous and heterozygous genomic regions, all resequenced haploids displayed complete homozygosity across the entire genome (Figure 1G). Haploid plants derived from *HUAXU-ee* lines exhibited markedly reduced plant height, smaller glumes, and complete sterility (Figure 1H and 1I), further confirming their haploid identity. Together, these findings demonstrate that egg cell-specific expression of *HUAXU* efficiently induces parthenogenesis and haploid formation in hybrid rice.

### Fully penetrant clonal reproduction in hybrid rice

Coupling engineered parthenogenesis with clonal gametogenesis represents a major approach for achieving synthetic apomixis in sexual plants.^12-14^ To assess whether *HUAXU* can be engineered to induce synthetic apomixis, we constructed a vector, *HUAXU-ee_sgMiMe* (Figure 2A), by integrating the *HUAXU-ee* cassette into the *sgMiMe* vector, which has been designed to generate clonal gametes through the simultaneous knockout of *OSD1, PAIR1*, and *REC8*, as previously described.^16,17^ We then transformed this construct into the hybrid rice CY84 (WT) via *Agrobacterium*-mediated callus transformation. Through the high-throughput tracking of mutations (Hi-TOM) identification,^49^ we obtained five mutant lines for *HUAXU-ee_sgMiMe*, with all three *MiMe*-associated genes (*OSD1, PAIR1*, and *REC8*) exhibiting homozygous or biallelic frameshift mutations (Figure S7A). After verifying the presence of the egg cell-specific expression fragment of *HUAXU-ee* in all five lines from *HUAXU-ee_sgMiMe* (Figure S7B), we designated these lines as *Fixation of Hybrids 8* (*Fix8)* #1 to #5 hereafter. Compared to the WT, the T_0_ plants of these *Fix8* lines did not show any obvious differences in plant and panicle morphology (Figures 2B and 2C). The seed setting rates of *Fix8* T_0_ plants varied from 50.4±3.9% to 74.3±2.4% across the five *Fix8* lines, compared to the WT (74.9±2.4%) (Figures 2C and 2D). Notably, among the five plants, the *Fix8* T_0_ #3 plant showed nearly the same seed setting compared to the WT plants (Student’s *t*-test, two-tailed, P > 0.05) (Figures 2C and 2D).

**Fig. 2.**
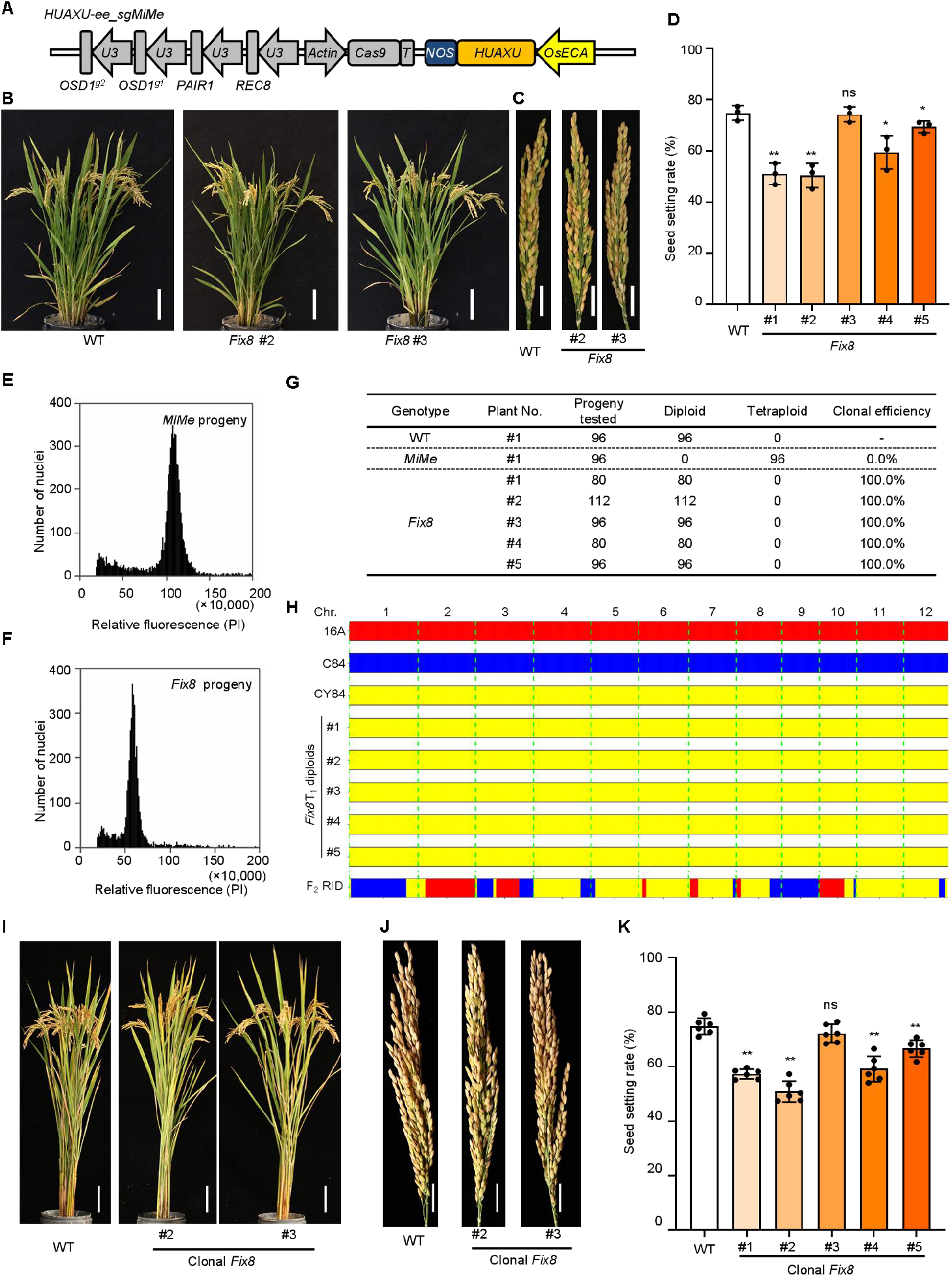
Full penetrance of apomictic hybrid rice induced by combining *MiMe* with egg cell–ectopic expression of *HUAXU*. (A) Schematic of the *HUAXU-ee_sgMiMe* construct, designed for egg cell-specific *HUAXU* expression and simultaneous knockout of *OSD1, PAIR1*, and *REC8* to induce *MiMe* and clonal gamete formation. Two sgRNAs targeted *OSD1* to enhance homozygous mutation efficiency. (B-C) Plant (B) and panicle (C) morphology of WT and *Fix8* T_0_ plants. Scale bars, 20 cm (B); 2 cm (C). (D) Seed setting rate of *Fix8* T_0_ plants compared to WT. Data represent the mean ± SD from three primary panicles per plant (n=3). **p < 0.01; *0.01 ≤ p < 0.05; ns, not significant (p ≥ 0.05; two-tailed Student’s *t*-test). (E–F) Ploidy determination of progeny from *Fix8* (E) and *MiMe* (F) lines by flow cytometry. PI, propidium iodide. (G) Ploidy analysis of the progeny from WT, *MiMe*, and *Fix8* lines. (H) Whole-genome sequencing of diploid *Fix8* T_0_ progeny. SNPs specific to parental alleles 16A (red), C84 (blue), or both (yellow) across the 12 rice chromosomes. *Fix8* diploid progeny exhibited genome-wide heterozygosity, closely matching the CY84 hybrid F_1_, whereas CY84 F_2_ recombinant inbred lines (RIDs) showed recombination and partial homozygosity. (I-J) Plant (I) and panicle (J) morphology of WT and clonal Fix8 progeny grown in paddy fields. Scale bars, 20 cm (I); 2 cm (J). (K) Seed setting rate of clonal *Fix8* T_1_ plants compared to WT. Data represent the mean ± SD from six plants (three primary panicles per plant). **p < 0.01; ns, not significant (p ≥ 0.05; two-tailed Student’s *t*-test).

Homozygous *MiMe* mutants are known to consistently produce tetraploid progeny due to genome duplication, whereas clonal reproduction typically yields diploid offspring.^15-17,20^ To determine whether *Fix8* T_0_ lines give rise to clonal progeny, we performed flow cytometric analysis on offspring from all five *Fix8* lines to assess their ploidy levels. To facilitate detection, a *MiMe* line was generated as a control by transforming the *sgMiMe* vector into the hybrid rice CY84 (Figure S7A). As expected, all 96 analyzed progeny from the *MiMe* line were tetraploid (Figures 2E, 2G, and S8), consistent with previous reports.^16,17^ In contrast, flow cytometric profiling of 80, 112, 96, 80, and 96 progeny from the five *Fix8* lines revealed that all offspring were diploid, with no tetraploid individuals detected (Figures 2F, 2G, and S8). To confirm that the diploid progeny of *Fix8* indeed resulted from clonal reproduction rather than chromosomal recombination, we genotyped offspring from all five *Fix8* lines using 12 InDel markers. The analysis revealed that although homozygous loci were detected in F_2_ RID plants as controls, all progeny from *Fix8* T_0_ plants consistently retained heterozygous genotypes similar to the WT hybrid CY84 at all 12 InDel loci (Figure S9). To further validate whether the *Fix8* progeny maintained heterozygosity across the entire genome like the hybrid CY84, we performed whole-genome sequencing with an average coverage depth of 20**×**. The results demonstrated that, unlike the F_2_ RID plant, which displayed evidence of chromosomal recombination and homozygous genomic regions, all *Fix8* progeny maintained complete heterozygosity. They faithfully inherited the full complement of hybrid genotypes from their maternal CY84 across the entire genome (Figure 2H). These results further confirm that the diploid progeny of *Fix8* T_0_ plants were indeed generated through clonal reproduction. Together, the consistent diploid ploidy and genotypes fully aligned with those of the hybrid F_1_ across all the detected progeny of *Fix8* T_0_ plants demonstrate that the diploid offspring of *Fix8* are clonally derived, with a clonal efficiency of 100% across all five obtained lines.

### Stable clonal reproduction and agronomic performance of *Fix8* hybrid rice across successive generations

Stable inheritance of seed set and clonal efficiency is essential for the application of synthetic apomixis. To determine whether the diploid *Fix8* progeny (hereafter termed clonal *Fix8*) retained the heterosis of the F_1_ hybrid, clonal *Fix8* plants in the T_1_ generation were cultivated under field conditions, and their agronomic traits were systematically evaluated. The results showed that except for one line that displayed a dwarf phenotype, possibly due to the position effect of T-DNA insertion, four independent lineages showed no significant differences in plant height, tiller number, and panicle length compared to the WT F_1_ hybrid CY84 (Figures 2I, 2J, and S10). Similar to their T_0_ parents, seed setting rates of T_1_ clonal *Fix8* plants ranged from 50.9 ± 3.5% to 72.2 ± 3.0% across independent lines, accounting for around 68.0% to 96.5% of F_1_ hybrid WT control (74.8 ± 2.7%) (Figures 2J and 2K). Notably, the *Fix8* #3 line also displayed a near-normal seed setting compared to the WT F_1_ plants (Student’s *t*-test, two-tailed, P > 0.05) (Figures 2J and 2K).

Since *Fix8* #3 line constantly exhibited plant morphology and seed set comparable to the WT F_1_ hybrid in both T_0_ and T_1_ generations, we next selected this line to test its potential for agricultural applications. We planted the T_2_ progeny of *Fix8* #3 in three larger populations derived from individual T_1_ plants to assess their clonal efficiency.Among 1104, 1180, and 806 T_2_ progeny tested, 1101, 1176, and 804 diploid individuals were identified (Figures 3A–3C), corresponding to clonal efficiencies of 99.7%, 99.7%, and 99.8%, respectively. These results demonstrate that *Fix8* represents a near-fully penetrant synthetic apomixis hybrid rice and indicate that the clonal reproductive traits of *Fix8* are stably inherited across successive generations.

**Fig. 3.**
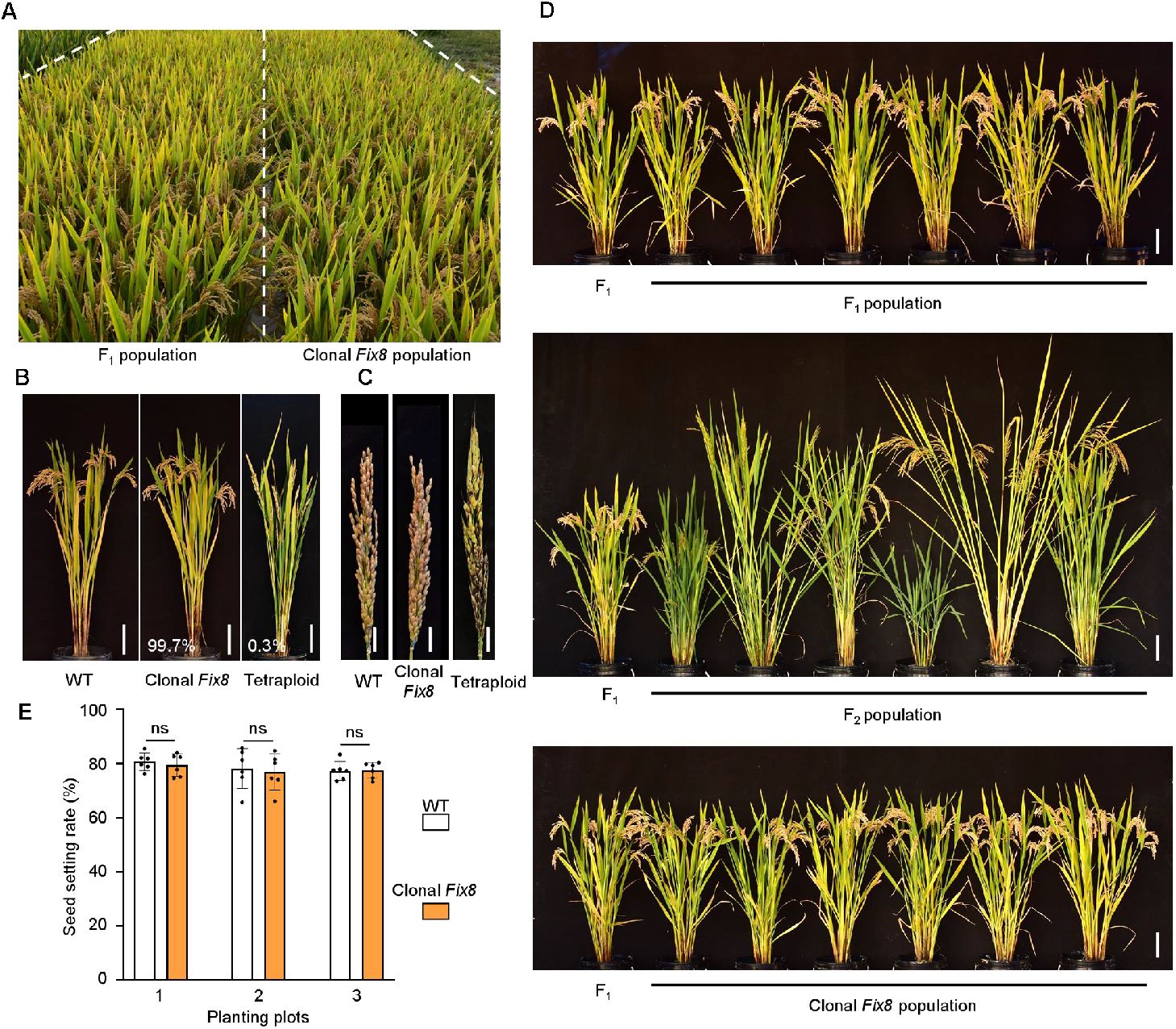
Stable inheritance of nearly complete penetrance of apomixis and normal seed set in *Fix8*. (A) Population performance of WT and clonal *Fix8* line #3 in the T_2_ generation. (B) Clonal diploid progeny and rare induced tetraploids from *Fix8* line #3 in the T_2_ generation. Scale bar: 20 cm. (C) Panicles of the progeny from *Fix8* line #3 in the T_2_ generation. Scale bar, 2 cm. (D) Morphological comparison of F_1_ (Hybrid CY84, WT), F_2_ (RIDs of CY84), and clonal *Fix8* T_2_ plants. Scale bar, 20 cm. (E) Seed setting rate of *Fix8* line #3 in the T_2_ generation versus WT grown in parallel plots. Data are mean ± SD from six randomly selected plants. ns, not significant (p ≥ 0.05; two-tailed Student’s *t*-test).

As the plant morphology and fertility are also highly associated with the growth conditions, to test whether the fertility is affected under different field environments, we finally planted the T_2_ progeny of #3 line and WT in parallel-planting different field plots, with varying soil properties, water availability, topography, and management practices. This comparison revealed that clonal *Fix8* T_2_ plants consistently exhibited plant and panicle morphology comparable to those of WT F_1_ hybrid plants (Figures 3A-3D). Remarkably, these clonal *Fix8* T_2_ plants exhibited uniform morphology both within the lineage and relative to the WT F_1_ hybrid population, without signs of phenotypic segregation typically seen in F_2_ progeny (Figure 3D). Importantly, clonal *Fix8* #3 T_2_ plants displayed seed setting rates of 76.8 ± 6.1% to 79.4 ± 3.8% under varying field conditions, which were consistently comparable to the WT (77.3 ± 3.2% to 80.7 ± 2.9%) grown in parallel plots (Figure 3E). Together, these findings confirm that *Fix8* #3 is a nearly fully penetrant apomictic hybrid rice capable of stable clonal reproduction, normal or near-normal seed production, and consistent agronomic performance across generations.

## DISCUSSION

Crop uniformity is a cornerstone of modern agriculture, essential for predictable cultivation management and yield stability. This necessity is reflected in stringent global seed purity standards; for example, in China, inbred rice must exceed 99% purity, and hybrid seeds must surpass 97%. A major hurdle for the application of synthetic apomixis has been the simultaneous achievement of high clonal efficiency and normal fertility necessary to meet these commercial benchmarks. In this study, we present a *HUAXU*-based synthetic apomixis system that overcomes this challenge. Our system achieves near-complete (>99%) clonal seed production across all independent lines and across multiple generations, even when evaluated in large populations. This efficiency not only meets but reliably exceeds current seed purity standards, demonstrating its immediate potential for commercial deployment. By enabling the indefinite fixation of heterosis, this technology offers a practical and cost-effective route to sustain hybrid vigor in agriculture, eliminating the need for recurrent and costly hybrid seed production.

A notable finding of our work is the superior performance of the sperm-cell-specific transcription factor *HUAXU* compared to previously reported parthenogenesis inducers, such as *OsBBM1, OsBBM4*, and *OsWUS*. While ectopic expression of broadly expressed transcription factors can trigger parthenogenesis, it often leads to significantly reduced seed set or compromised clonal efficiency. We propose that *HUAXU*, as a natural sperm-derived factor, may more precisely initiate the embryonic program in the egg cell, resulting in a more developmentally congruent and less disruptive process. This is supported by the consistent production of *Fix8* lines that exhibit not only over 99% clonal reproduction efficiency but also normal plant morphology and seed set approaching wild-type levels—a combination rarely achieved with previous systems. The high efficiency and stability of clonal reproduction in our *HUAXU*-based system, faithfully maintaining heterozygosity across the genome over generations, underscore its robustness and represent a significant advance towards usable synthetic apomixis.

While the *HUAXU*-based system achieves near-perfect clonal fidelity, the rare occurrence of tetraploid progeny warrants attention. These individuals can be readily identified and rogued in the field post-flowering due to their distinct morphology, ensuring seed lot purity during multiplication. Furthermore, as tetraploid plants are typically less fertile, their contribution to the seed bank would be naturally limited even without intervention. Nevertheless, achieving 100% penetrance for clonal diploid progeny remains a valuable objective for future research, potentially through further optimization of the expression cassette or the identification of even more specific embryogenesis triggers.

In conclusion, the *HUAXU*-based synthetic apomixis system described here constitutes a pivotal milestone in seed technology. It provides a robust, reproducible, and agronomically viable strategy to permanently fix heterosis in hybrid crops through seeds. This breakthrough paves the way for transformative changes in breeding paradigms and seed production systems, promising a new era of sustainable agricultural productivity.

## ACKNOWLEDGMENTS

We thank Liping Xu, Xingchen Jin, Zhengyuan Hong, Rongwei Cui, Weilong Yuan, Qing Liu, Fuchun Zhang, and Yuchun Rao (Zhejiang Normal University) for assistance with rice cultivation and data collection. We also acknowledge the Public Laboratory of the China National Rice Research Institute for technical support with the Zeiss LSM 980 confocal microscope. This work was supported by the National Natural Science Foundation of China (32025028 and 32188102), the National Key Research and Development Program of China (2022YFF1003304), the earmarked fund for CARS (CARS-01-10), and the Agricultural Science and Technology Innovation Program (CAAS-ASTIP-2021-CNRRI).

